# Mechanism of the Switch from NO to H_2_O_2_ in Endothelium-Dependent Vasodilation in Diabetes

**DOI:** 10.1101/2021.05.18.444667

**Authors:** Cody Juguilon, Zhiyuan Wang, Yang Wang, Anurag Jamaiyar, Yanyong Xu, Molly Enrick, James Gadd, Chwen-Lih W. Chen, Autumn Pu, Chris Kolz, Vahagn Ohanyan, Yeong-Renn Chen, Yanqiao Zhang, William M Chilian, Liya Yin

## Abstract

Coronary microvascular dysfunction is prevalent among diabetics and is correlated with cardiac mortality. Compromised endothelial-dependent dilation (EDD) is an early event in the progression of diabetes, but the mechanisms remain incompletely understood. Nitric oxide (NO) is the major endothelium-dependent vasodilatory metabolite in the healthy coronary circulation, but switches to hydrogen peroxide (H_2_O_2_) in coronary artery disease (CAD) patients. Because diabetes is a major risk factor for CAD we hypothesized that a similar switch from NO-to-H_2_O_2_ occurs in diabetes. Methods: Vasodilation was measured *ex vivo* in isolated coronary arteries from wild type (WT) and microRNA-21 (miR-21) null mice fed chow or high fat and sugar diet, and LepR null (db/db) mice using myography. Myocardial blood flow (MBF), blood pressure, and heart rate were measured *in vivo* using contrast echocardiography and a solid-state pressure sensor catheter. RNA from coronary arteries, endothelial cells and hearts were analyzed via qPCR for gene expression and protein expression was assessed via Western-Blot analyses. Superoxide was detected via electron paramagnetic resonance (EPR). Results: 1) *Ex vivo* coronary EDD and *in vivo* MBF was impaired in diabetes. 2) L-NAME (NO-synthase inhibitor) inhibited *ex vivo* coronary EDD and *in vivo* MBF in WT, while PEG-catalase (H_2_O_2_ scavenger) inhibited diabetic EDD *ex vivo* and MBF *in vivo.* 5) miR-21 deficiency blocked the NO-to-H_2_O_2_ switch and prevented diabetic vasodilation impairments. 6) Diabetic mice displayed increased serum NO and H_2_O_2_, upregulated mRNA expression of Sod1, Sod2, iNos, and Cav-1, and downregulated Pgc-1α. Deficiency of miR-21 reversed these changes. 7) miR-21 deficiency increased PGC1α, PPARα and eNOS protein and reduced detection of endothelial superoxide. Conclusions: Diabetics exhibit an NO-to-H_2_O_2_ switch in the mediator of EDD coronary dilation, which contributed to microvascular dysfunction and is mediated by miR-21. This study represents the first mouse model recapitulating the NO-to-H_2_O_2_ switch seen in CAD patients.

## Introduction

CAD is a leading cause of mortality and morbidity in the United States. Although the principal focus of CAD research is on obstructive epicardial disease, there is increasing awareness of coronary microvascular dysfunction within the context of ischemic heart disease [1]. Coronary microvascular dysfunction is prevalent in diabetic patients and is independently correlated with cardiac mortality among both diabetics and non-diabetics [26, 43]. Metabolic disorders, especially diabetes mellitus, increases the morbidity and mortality of many microvascular complications, such as diabetic retinopathy [3, 36], diabetic nephropathy [8] and diabetic cardiomyopathy. The mortality of patients with CAD and diabetes mellitus increases 2- to 4-fold compared to those nondiabetic patients with CAD [3]. Diabetic patients have a higher risk of developing CAD and metabolic disorders like hyperglycemia and dyslipidemia that could contribute towards endothelial dysfunction by reducing bioavailability of NO and increasing the level of reactive oxygen species (ROS) [30, 47].

Nitric oxide (NO) is the major endothelium-derived relaxing factor (EDRF) and contributes to the endothelium-dependent dilation (EDD) in coronary arteries under normal conditions [3, 13]. Previous work by Dr. Gutterman’s group has shown that the mediator of flow-induced vasodilation in patients with CAD switches from NO-dependent to H_2_O_2_-dependent vasodilation, while no such switch occurs in aged patients without CAD [7, 26],[3, 6, 28]. Despite the fact that both NO and H_2_O_2_ are vasodilators, they each have their own inherent properties [3, 62]. H_2_O_2_ plays important roles in pathological conditions such as atherosclerosis and hypertension where it is involved in compensation of vasorelaxation in large vessels [33]. This NO-to-H_2_O_2_ transition in EDD might be an early pathologic step in the progression of CAD [37, 41, 50].

Increased ROS levels and reduced NO bioavailability play a key role in the induction and progression of microvascular and cardiovascular complications during diabetes. It is well documented that in diabetic animals, EDD in coronary arterioles is impaired [4, 12, 48]. However, the underlying mechanism of such impairment is not completely understood. The overlap of microvascular dysfunction in both CAD and diabetic vascular diseases prompted us to explore whether the NO-to-H_2_O_2_ switch mediating vasodilation reported in CAD patients also occurs in diabetes, as well as the underlying mechanism of such a switch. In this study, we tested this hypothesis in diabetic mouse models, focusing on aortic (macro) and coronary (micro) vasodilation.

To address the mechanism of the switch between NO and H_2_O_2_, we focused on a specific microRNA that we previously found upregulated in diabetes. It is important to note that diabetes is associated with increased expression of microRNAs (miRNAs), which are short non-coding RNA molecules that bind to the 3’ UTRs of target genes and regulate gene expression transcriptionally and post-transcriptionally. Micro-RNA 21 (miR-21) plays important roles in metabolism [26, 66],[9, 51] and is enriched in endothelial cells (EC) where it regulates NO production, cell proliferation and apoptosis [61, 67], and modulates vascular diseases and remodeling [57, 59]. In this study, we investigated if miR-21 plays any role in the NO-to-H_2_O_2_ switch in the mediator of EDD in diabetes using a genetic knockout model.

## Material and Methods

### Animals

All procedures were conducted with the approval of the Institutional Animal Care and Use Committee of the Northeastern Ohio Medical University and in accordance with National Institutes of Health Guidelines for the Care and Use of Laboratory Animals (NIH publication no. 85-23, revised 1996). Mice were housed in a temperature-controlled room with a 12:12-hour light-dark cycle and maintained with access to food and water *ad libitum.* C57BL/6J wild type (WT) mice, B6.BKS(D)-Leprdb/J (db/db) mice, and miR-21^-/-^ mice were purchased from the Jackson Laboratories. The miR-21^-/-^ mice from JAX were of a mixed background with several backcrosses with C57BL/6NJ and backcrosses were continued for at least 6 more generations onto the C57BL/6NJ background. Littermate WT and miR-21^-/-^ mice fed chow diet were used as controls. For diet-induced diabetic mice (WT+ HFHS or miR-21^-/-^+ HFHS), 6-week-old mice were fed high fat and high sugar (HFHS) diet (Envigo #TD.88137 with 42.7% carbohydrate [mostly sucrose], 42% fat) for at least 5 months [55].

### Cell culture

Different lot numbers of WT and db/db mouse coronary endothelial cells (WT and db/db mCEC) were purchased from Cell Biologics; healthy human and diabetic patient coronary artery endothelial cells (Healthy and Diabetic hCEC) were purchased from Lonza. The cells were cultured in the medium supplied from the companies as instructed. Cells less than passage six were used in the experiments. For low and high glucose treatment, cells were cultured in media supplemented with 5.5mM (LG) and 25.5mM (HG) glucose for 72 hours. For high lipid (HL) treatment, cells were cultured supplemented with 150μM Linoleic Acid, Oleic Acid, and Cholesterol for 72 hours with 1% serum.

### Isolation of mouse coronary endothelial cells (mCEC)

mCEC were isolated with the modified protocol from <u>Vascular Research – Dept. of Pathology – BWH (harvard.edu)</u> and the publication [38]. Briefly, the mouse heart was dissected and minced into small pieces. After the digestion of the heart using collagenase I (Worthington), cells were washed and incubated with Dynabeads conjugated with anti-CD31 antibody (Thermofisher). The beads with EC were washed several times and cultured in mouse EC cultured medium (Cell Biologics). When the cells were confluent, they were purified again with Dynabeads conjugated with anti-Mouse CD102 (ICAM2) antibody. Cells less than six passages were used for the experiments.

### Isolated vessel studies

The thoracic aorta was isolated and aortic segments were excised, cannulated, and incubated in PSS buffer (145mM NaCl, 4.7mM KCl, 2.0mM CaCl_2_·2H_2_O, 1.17mM MgSO_4_·7H_2_O, 3.0mM MOPS, 1.2mM NaH_2_PO_4_·H_2_O, 5.0mM Glucose, 2.0mM Pyruvic Acid, 0.02mM EDTA, PH7.38-7.40) at 37°C in the bath chamber of Multimyograph system 620M (DMT, Danish Myo Technology A/S). Small coronary arteries (100-200 μm internal diameter) were dissected from the epicardial surface of the left ventricle under a dissecting microscope. After these preparations, two small (20 μm diameter) wires were inserted into the coronary arteries and connected to a force transducer. After optimization of tension, coronary arteries were precontracted with the thromboxane mimetic U46619 (1μM). Once a stable contraction was achieved, pharmacological agents were administered in the bath and cumulative concentration-response curves were obtained. Endothelium-dependent dilation (EDD) was measured by acetylcholine (Ach), and endothelium-independent dilation was measured by sodium nitroprusside (SNP). In some studies, pharmacological inhibitors were administered into the bath chamber (100μM/L N^G^-nitro-L-arginine-methyl ester [L-NAME], 500unit/ml PEG-catalase, 10μM/L indomethacin) for 30 min before recording the vasodilation of mouse arteries.

### Nitrite and nitrate measurement

The serum levels of nitrite plus nitrate (NOx) were assessed. NOx was measured by a commercial kit (Abcam) according to the manufacturer’s instructions. Briefly, serum was diluted and ultra-filtered through a 10 kDa cutoff filter. The nitrate was converted to nitrite with nitrate reductase based on the Griess reaction. NOx concentrations were determined at OD 540 nm with a SpectraMax Plus microplate spectrophotometer (Molecular Devices).

### Measurement of H_2_O_2_

The levels of H_2_O_2_ were measured with a H_2_O_2_ assay kit (Abcam #ab65328) according to the manufacturer’s instructions. Briefly, the working solution containing OxiRed probe and horse radish peroxidase (HRP) was added to serum samples. In the presence of HRP, the OxiRed probe reacts with H_2_O_2_ to produce a product that can be detected at OD570 nm by a SpectraMax Plus microplate spectrophotometer (Molecular Devices).

### Superoxide measurement by X-band EPR (electron paramagnetic resonance)

EPR spintrapping measurements of superoxide from coronary arterial ECs were performed with a Bruker EMX Micro spectrometer operating at 9.43 GHz, and the spin trap 5,5 dimethyl-1-pyrroline-N-oxide (DMPO) at a final concentration of 50 mM was used. Care was taken to keep the DMPO-containing solutions covered to prevent any light-induced degradation. The DMPO (ultrahigh purity) was purchased from Dojindo (Rockville, MD). The cell culture containing DMPO (50 mM) was transferred to a 50 ml capillary tube (Drummond Wiretrol, Broomall, PA) loaded into the EPR resonator (HS cavity, Bruker Instrument, Billerica, MA). EPR spectra were recorded at room temperature with the following parameters: center field 3360 G, sweep width 100 G, power 20 mW, receiver gain 1 × 10^5^, modulation amplitude 1 G, conversion time 83 ms, time constant 327.68 ms, and number of scans 12. Measurements were performed at X-band with 100-kHz modulation frequency with 10-mW microwave power, and a modulation amplitude of 1.0 Gauss using an EMX HS resonator. The spectral simulations for SOD-sensitive DMPO/’·OH adducts spin quantitation were performed using the WinSim program developed at NIEHS by Duling [20] using the hyperfine coupling constants a^N^=a_β_^H^=14.88 G. The intensity of EPR signal was presented by spin number calculated from double integration of simulated spectrum [29].

### RNA isolation and qPCR

RNA was extracted from tissues and cells using Trizol Reagent (Thermo Fisher) or RNeasy Mini Kit (Qiagen) and mRNA levels of gene expression were quantified by quantitative real-time PCR (qRT-PCR) using SYBR Green (GeneCopoeia) on a 7500 Real-Time PCR machine (Applied Biosystems). mRNA levels were normalized to 36B4 and calculated the relative fold change compared to the controls. Primers were designed and synthesized by IDT (Integrated DNA Technologies); sequences are listed in Table 1. MicroRNAs were extracted from cells and tissues using miRNeasy Mini Kit (Qiagen). Reverse transcription was done with TaqMan MicroRNA Reverse Transcription Kit and quantified using a TaqMan MicroRNA assay kit and TaqMan Universal Master Mix II, no UNG (Thermo Fisher) for real-time-PCR, then normalized to U6 and fold change was calculated and compared to the control as described [65].

### Western blot

Protein was extracted from mouse hearts, quantitated, run on the 12% pre-cast mini gel (BioRad), transferred and blotted with Peroxisome proliferator-activated receptor alpha (PPARα, ab 215270), endothelial nitric oxide synthase (eNOS, sc-376751) and Glyceraldehyde 3-phosphate dehydrogenase (GAPDH, Thermo Fisher, #10941-1-AP) antibodies as described [65]. The protein expression level is normalized to GAPDH.

### Hemodynamics and myocardial blood flow measurements and calculation of Cardiac Work

Heart rate (HR), mean arterial pressure (MAP), and myocardial blood flow (MBF) were measured as previously described [44, 45]. Briefly, mice were anesthetized, the right jugular vein was cannulated for intravenous (i.v.) drug and contrast echocardiography reagent infusions, the femoral artery was cannulated with a 1.2F pressure catheter (Scisense Inc, Ontario, Canada) to measure MAP and HR. Cardiac work (CW) directly reflects myocardial oxygen consumption, so we employed this index as a surrogate for oxygen consumption to decipher the basis of coronary metabolic dilation. Metabolic demands in the heart are calculated by the double product (DP) of HR and MAP. Double product is expressed as CW in the figures and is calculated by the formula CW= HR x MAP. MBF was measured from 3-5 different images obtained from the same condition (baseline and treatments with norepinephrine [2.5μg/kg/min and 5μg/kg/min], L-NAME [100mg/kg] or PEG-Catalase [100^u^/g] after hexamethonium [5mg/kg]).

### Statistical Analysis

Data are presented as mean ± the standard error of the mean (SEM). Data were analyzed using SPSS19.0 and PRISM 5.0 statistical software. Comparisons between groups were made using one-way analysis of variance (ANOVA) followed by Bonferroni post-hoc test. Comparisons between two groups were made using unpaired student t-test. Differences were considered statistically significant at a value of *P* < 0.05. The test of normal distribution shows that the data distributions were not significantly different from normal.

## Results

### NO-mediated Ach-induced EDD in aortic arteries from control and diabetic mice

Thoracic aortic arteries from 9-month-old WT, db/db and diet-induced diabetic (WT+HFHS) mice were isolated for vasodilation induced by Ach. The plasma lipid and glucose levels in WT mice on HFHS diet was significantly increased than the WT mice on chow diet. (Supplementary Figure 1A). The glucose tolerance test (GTT) showed that WT mice on HFHS exhibited a diabetic phenotype (Supplementary Figure 1B). Compared to WT mice, Ach-induced EDD was decreased in thoracic aortic arteries from db/db mice and WT+HFHS mice (Figure 1A). Ach-induced EDD was inhibited in all groups after treatment with the NO synthase inhibitor L-NAME. Figure 1B. illustrates that the concentration-dependent relaxation responses to SNP was equivalent in all the three groups. These data suggest NO is the mediator of Ach-induced EDD in aortic arteries from both WT and diabetic animals.

**Fig 1.**
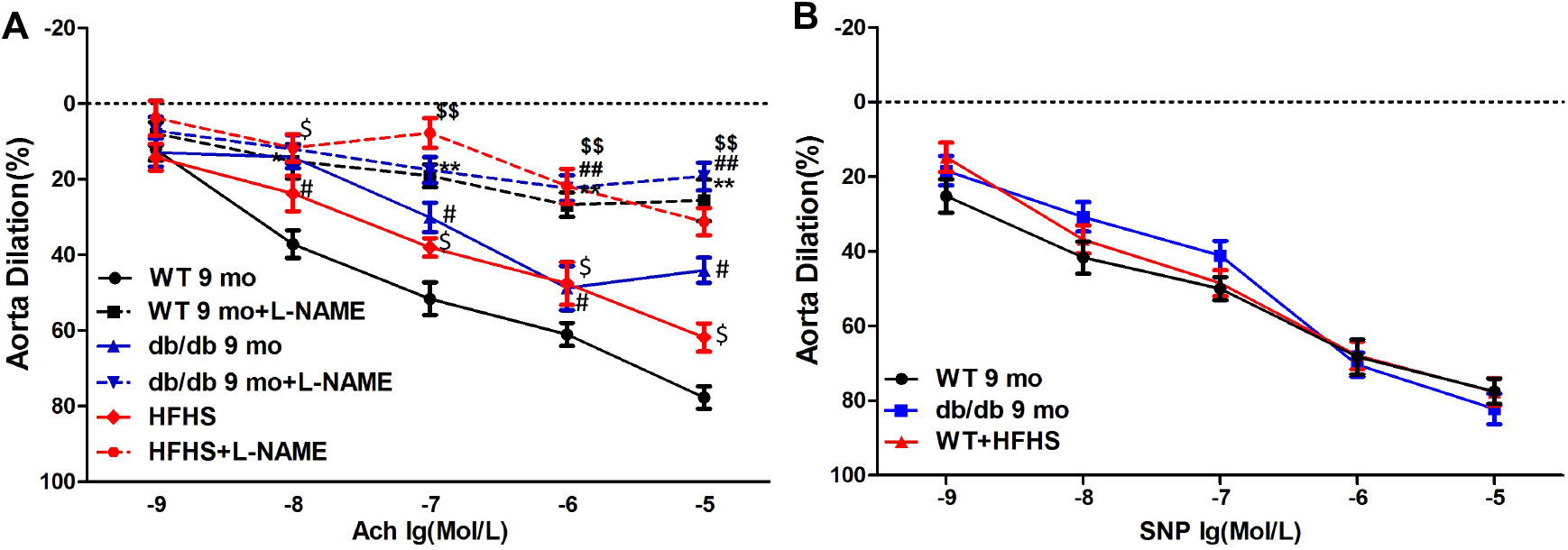
Vasodilation was mediated by NO in aortic arteries from both WT and diabetic mice. (A) Compared with WT, Ach-induced EDD was decreased in aortic arteries from both 9-month-old db/db and diet-induced diabetic (WT+HFHS) mice. Additionally, L-NAME significantly inhibited Ach-induced EDD in all aortic arteries. \**P*<0.05, \*\**P*<0.01 *vs.* WT, ^#^*P*<0.05, ^##^*P*<0.01 *vs.* db/db and ^$$^*P*<0.01 *vs*. WT+HFHS. (B) SNP-induced aorta vasorelaxation was unchanged in WT, db/db and WT+HFHS mice. (n=6).

### The mediator of Ach-induced EDD in diabetic coronary arteries switches from NO to H_2_O_2_

To determine if NO is also the mediator of Ach-induced EDD in the healthy coronary microcirculation, coronary arteries from 3-month, 9-month and 32-month-old mice were isolated for vasodilation studies. Figure 2A, 2B, and 2C show L-NAME inhibited Ach-induced EDD in coronary arteries from 3-month, 9-month, and 32-month-old WT mice, respectively. To compare the roles of NO to other endothelium-derived mediators of vasodilation, we also tested the impact of the H_2_O_2_ scavenger, polyethylene glycol-catalase (PEG-catalase) and the prostaglandins (PGs) synthesis inhibitor indomethacin on Ach-induced EDD in the coronary arteries. Figure 2A-C show that neither PEG-catalase nor Indomethacin had a significant effect on Ach-induced EDD in the coronary arteries in WT mice of all ages. The relative contributions of three endothelium-derived relaxing factors (PGs, NO, and H_2_O_2_) to Ach-induced EDD is summarized in Figure 2E. In coronary arteries from young and old WT mice, Ach-induced EDD was mostly mediated by NO. This suggests that NO plays a dominant role in regulation of Ach-induced EDD in WT mice at all ages. Furthermore, endothelium-independent relaxation is illustrated in Figure 2D, showing that the vascular smooth muscle cell response to SNP was not significantly changed in all the three groups.

**Fig 2.**
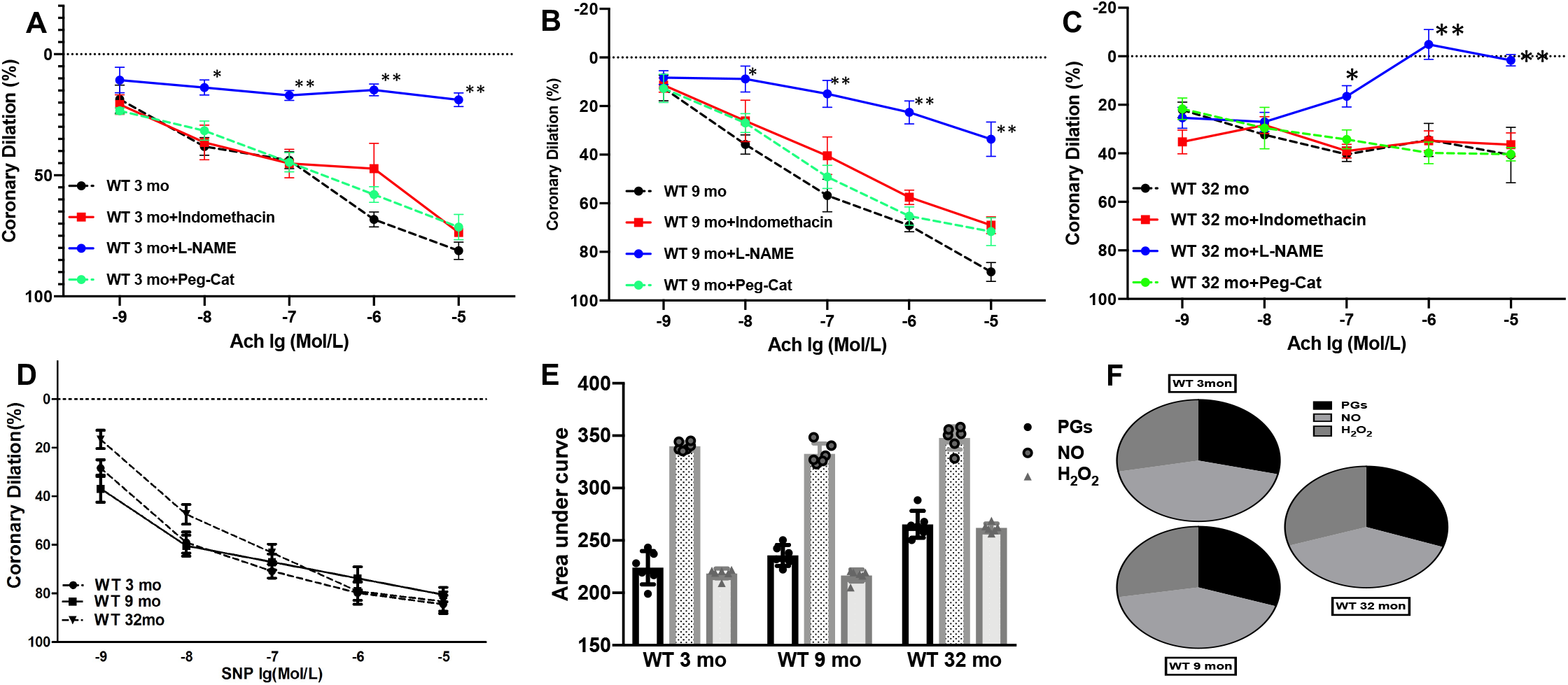
Coronary vasorelaxation in WT mice at different ages was NO-dependent. (A-C) L-NAME significantly inhibited Ach-induced EDD in 3-month, 9-month and 32-month-old WT mice respectively, but Peg-cat and Indomethacin had no significant effect on coronary dilation in WT mice. \**p*<0.05, \*\**p*<0.01 *vs.* WT 3-month in panel (A), \**p*<0.05, \*\**p*<0.01 *vs*. WT 9-month in panel (B), \**p*<0.05, \*\**p*<0.01 *vs.* WT 32-month in panel (C). (D) There was no difference in vasodilation in the small coronaries from WT mice at different ages after the treatment of SNP. (E and F) The relative contribution of vasodilators (PGs, NO and H_2_O_2_) to Ach-induced coronary dilation in WT mice was measured with area under curve and it shows NO plays a dominant role in coronary dilation in WT mice at different ages. n=6 mice/group.

After observing NO-mediated Ach-induced EDD in all coronary arteries of different ages of WT mice, we investigated whether NO is the major mediator of Ach-induced EDD in coronary arteries in diabetic mice. Figure 3A and 3B show that Ach-induced EDD was decreased in coronary arteries from 3-month and 9-month-old db/db mice. Importantly, L-NAME and indomethacin did not inhibit the Ach-induced EDD in coronary arteries from db/db mice. However, PEG-catalase inhibited the Ach-induced EDD in the diabetic coronary arteries. Figure 3C shows that HFHS diet induced diabetic mice (WT + HFHS) had decreased Ach-induced EDD compared to the control WT mice on chow diet. Furthermore, PEG-catalase, but not L-NAME or indomethacin, inhibited the Ach-induced EDD in these diet-induced diabetic mice. The results show that diet-induced diabetic mice phenocopied the db/db mice in vasodilation. Figure 3E and 3F showed the relative contributions of vasodilators (PGs, NO and H_2_O_2_) on the Ach-induced EDD in diabetic coronary arteries, suggesting that H_2_O_2_, instead of NO, plays the dominant role in Ach-induced EDD in diabetic coronaries. Figure 3D shows that no significant differences in SNP-induced coronary dilation were observed among all three groups.

**Fig 3.**
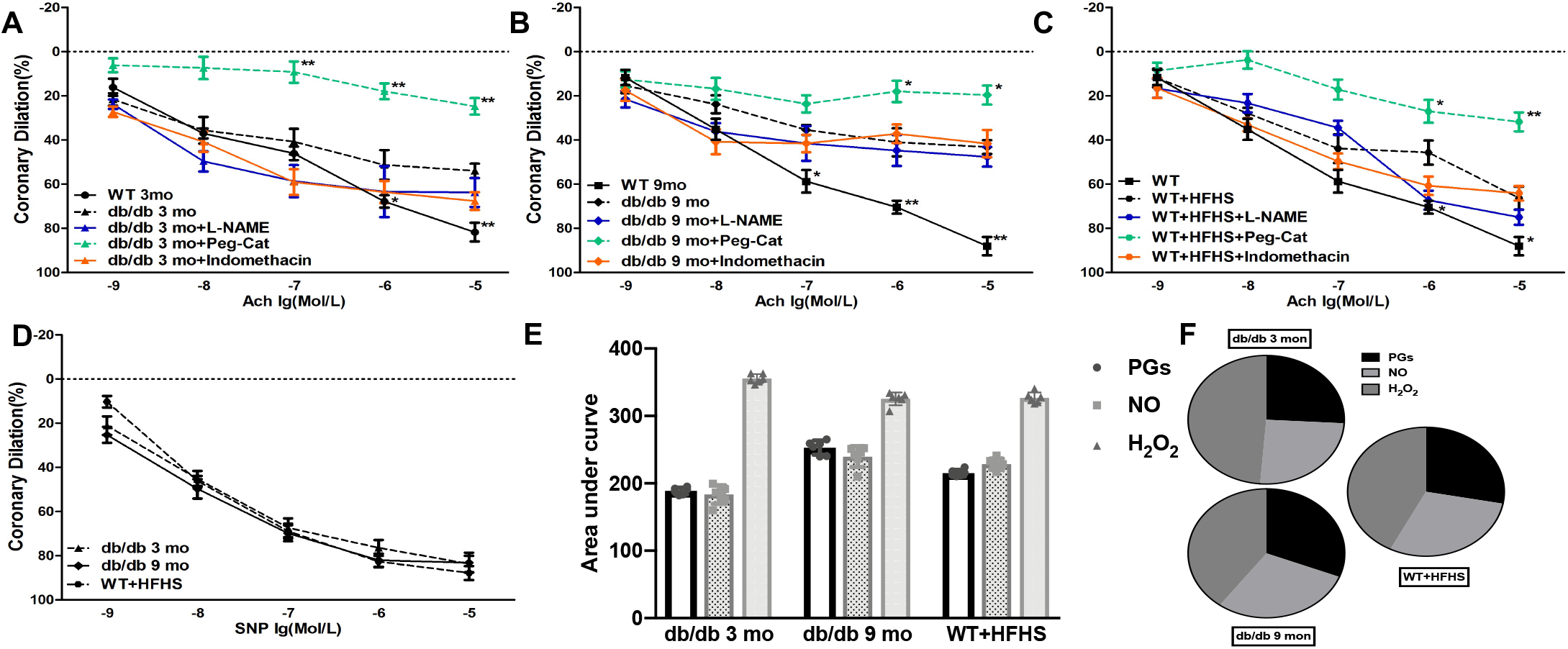
The mediator of Ach-induced EDD switches from NO to H_2_O_2_ in small coronary arteries from db/db and WT+HFHS mice. (A-C) Ach-induced EDD in coronary arteries from 3-month and 9-month-old db/db and 6-8-month-old WT+HFHS mice was significantly decreased compared to their WT age-matched controls. Peg-cat inhibited Ach-induced coronary dilation in db/db and WT+HFHS mice, but L-NAME and Indomethacin did not. \**p*<0.05, \*\**p*<0.01 *vs.* db/db 3-month in panel (A), \**p*<0.05, \*\**p*<0.01 *vs*. db/db 9-month (B), \**p*<0.05, \*\**p*<0.01 *vs.* WT +HFHS (C). (D) SNP-induced coronary vasorelaxation was unchanged in db/db and WT+HFHS mice. (E and F) The relative contribution of vasodilators (PGs, NO and H_2_O_2_) to Ach-induced coronary dilation in db/db and WT+HFHS mice was measured by area under curve. It suggests H_2_O_2_ plays a dominant role in coronary dilation in db/db and WT+HFHS mice. n=6 mice/group.

### Micro RNA-21 expression

miR-21 plays important roles in metabolism and cardiovascular diseases. To map which tissue expressed miR-21, we used our lacZ-miR-21 knockin mouse model to show miR-21 expression by ß-gal staining. Supplementary Figure 2 shows that miR-21 is highly expressed in the vessels in the heart, aorta, arterioles in skeletal muscles, but this microRNA was not abundantly expressed in other cell types of the heart. Figure 4A shows that compared to WT mice, miR-21 gene expression was significantly upregulated in the hearts and aortas of db/db mice. Figure 4B shows that both Healthy hCEC and Diabetic hCEC expressed significantly higher levels of miR-21 under HG and HL treatment. These data indicated that miR-21 expression is upregulated in diabetic hearts and CEC stimulated with HG and HL, suggesting that miR-21 plays important roles in diabetic cardiovascular dysfunction.

**Fig 4.**
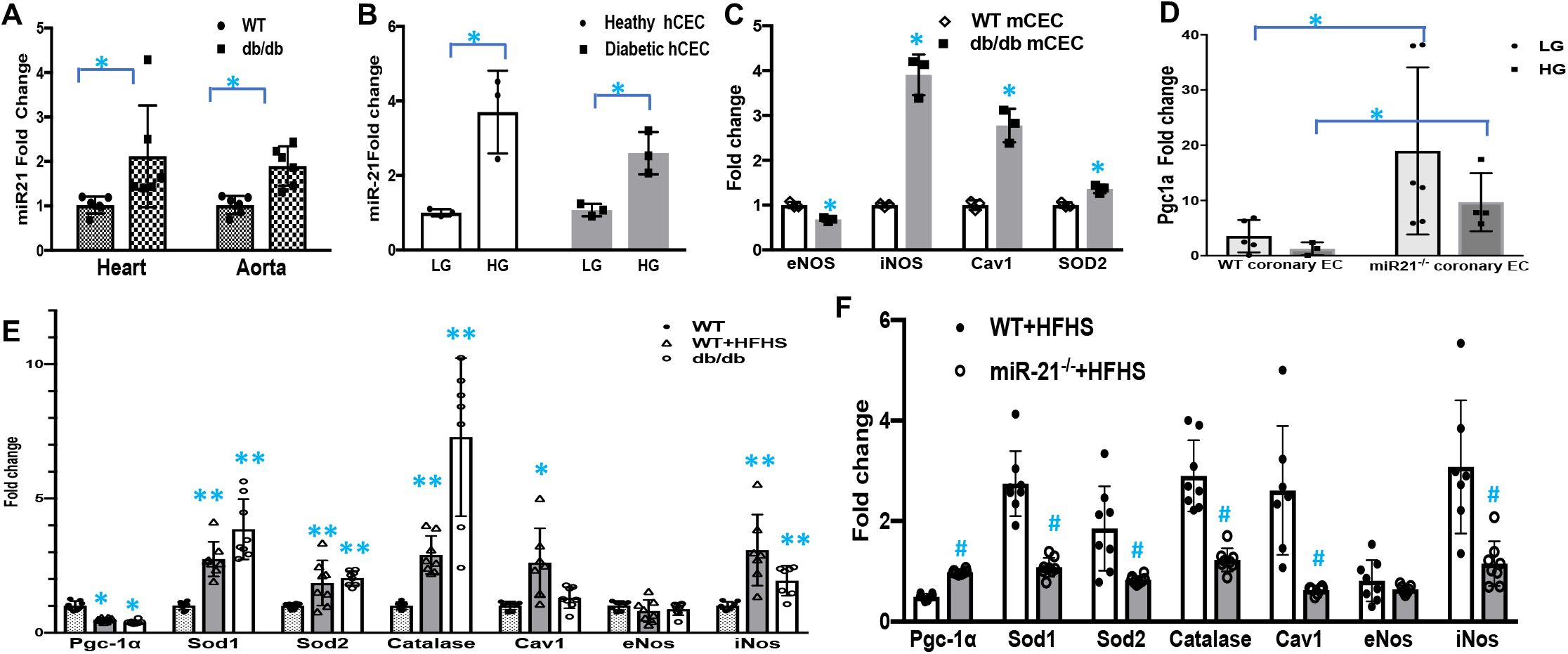
Gene expression by qPCR. A shows mi-R 21 expression in heart, aorta from WT and db/db mice (n=3, \**p*<0.05). (B) mi-R 21 expression was increased in healthy human coronary EC (hCEC) and diabetic human coronary EC (hCEC) under the treatment of high glucose (HG, 25.5mM glucose) compared to the cells under the treatment of low glucose (LG, 5mM glucose) (n=3, \**p*<0.05). (C) Compared to WT mouse coronary EC (WT mCEC), coronary endothelial cells in db/db mice (CEC) expressed significantly less eNos, higher iNos, Cav-1 and Sod2 (n=3, \**p*<0.05). (D) Expression of Pgc1α in coronary endothelial cells in miR-21^-/-^ was higher than in WTCEC subjected to LG and HG treatment. (*p<0.05, n=6). (E). Compared to WT mouse coronaries, expression of Sod1, Sod2, Catalase, Caveolin-1 and iNOS expression in coronaries increased; while Pgc-1α decreased in WT+HFHS and db/db mice (\**p*<0.05, \*\**p*<0.01 *vs*. WT). eNos expression did not change. (F) Gene expression in coronary arteries from WT+HFHS and miR-21^-/-^ +HFHS mice. Deficiency of miR-21 reversed the changes of gene expression in WT+HFHS mice (^#^*p*<0.05 *vs*. WT+HFHS, n=6 mice/group).

### Deficiency of miR-21 stopped the switch of the mediator of Ach-induced EDD from H_2_O_2_ to NO in coronary arteries of diabetic mice

Figure 5A shows that L-NAME significantly inhibited Ach-induced EDD in coronary arteries from miR-21^-/-^ mice on chow diet (miR-21^-/-^ + chow), but Peg-catalase and Indomethacin had no significant effect. This suggests that NO is the mediator of Ach-induced EDD in coronaries from miR-21^-/-^ mice on chow diet. Figure 5B shows that compared to WT mice on HFHS diet (WT + HFHS), Ach-induced EDD in coronaries was improved in miR-21^-/-^ mice on HFHS diet (miR-21^-/-^ + HFHS). Interestingly, L-NAME inhibited the Ach-induced EDD in coronaries of miR-21^-/-^ mice fed HFHS diet, but PEG-catalase and indomethacin did not. These data suggest that NO is the dominant mediator of Ach-induced EDD in coronary arteries from miR-21^-/-^ mice on HFHS diet, like that of the WT or the miR-21^-/-^ mice on chow. Figure 5C shows that no significant difference in SNP-induced coronary dilation among the groups, suggesting smooth muscle responsiveness to NO is unchanged. Figure 5D and 5E show the relative contributions of vasodilators (PGs, NO, and H_2_O_2_) to Ach-induced EDD in coronary arteries of the three groups, and suggest that NO, rather than H_2_O_2_, plays the dominant role in Ach-induced EDD in miR-21^-/-^ mice fed HFHS diet; therefore, deficiency of miR-21 blocked the NO-to-H_2_O_2_ switch in coronary arteries of diet-induced diabetic mice.

**Fig 5.**
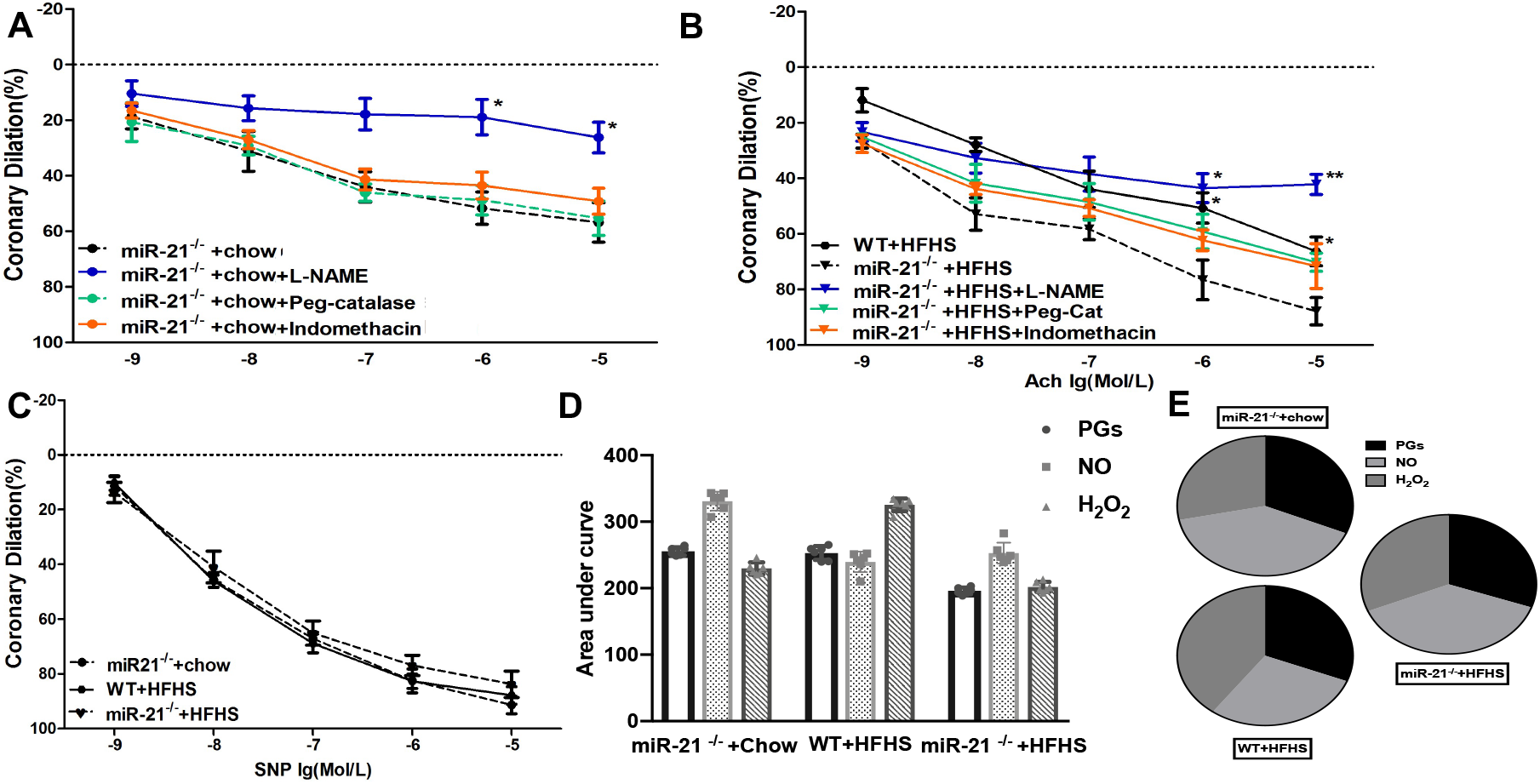
Deficiency of miR-21 reversed the switch in the mediator of Ach-induced EDD from H_2_O_2_ to NO in coronaries of WT+HFHS mice. (A) L-NAME significantly inhibited Ach-induced EDD in coronary arteries from miR-21^-/-^ on chow diet mice (miR-21^-/-^ + chow, but Peg-cat and Indomethacin had no significant effect on coronary dilation. \**p*<0.05 *vs.* miR-21^-/-^ +chow. (B) The vascular dilation to Ach was improved in coronaries from diet-induced diabetic mice from knockout background (miR-21^-/-^ +HFHS). L-NAME significantly inhibited Ach-induced EDD in miR-21^-/-^ +HFHS mice, whereas Peg-cat and indomethacin had no significant effect. \**p*<0.05, \*\**p*<0.01 *vs.* miR-21^-/-^ +HFHS in fig. 2C. (C) There was no difference in coronary dilation induced by SNP among coronaries in all mice. (D and E) The relative contribution of vasodilators (PGs, NO and H_2_O_2_) to Ach-induced EDD in miR-21^-/-^ +chow diet, WT+HFHS and miR-21^-/-^ +HFHS mice were measured with area under curve. n=6 mice/group.

### MicroRNA-21 regulated gene expression of vasodilation and ROS signaling pathways in diabetes

To investigate the underlying mechanism of the NO-to-H_2_O_2_ switch during the Ach-induced EDD in coronary arteries of diabetic mice, we used real-time RT-PCR to profile coronary arteries and mCEC from WT and diabetic mice to determine the genes that regulate the mediator transition from NO to H_2_O_2_ in Ach-induced EDD in diabetes. Figure 4C shows that compared to WT mCEC, db/db mCEC expressed significantly less eNos, but enhanced expression of inducible NO synthase (iNos), Caveolin-1 (Cav-1), and Superoxide dismutase 2 (Sod2), *p< 0.05, n=3, three different lot numbers of cells. Figure 4E shows the gene expression in the mouse coronary arteries, when compared to WT mice, gene expression of Sod1, Sod2, Catalase, Cav-1, and iNos was significantly upregulated in WT mice on HFHS diet and db/db mice, but eNos was not significantly changed (\**P*<0.05, \*\**P*<0.01 *vs.* WT, n=6). However, compared to WT, expression of PPAR-gamma co-activator-1 (Pgc-1α) was significantly reduced in WT mice on HFHS diet and db/db mice (\**P*<0.05 *vs.* WT, n=6). To investigate the effects of miR-21 on these genes, we also compared the gene expression of coronary arteries from miR-21^-/-^ mice fed HFHS diet and WT mice fed HFHS. Figure 4F showed that coronary arteries from miR-21^-/-^mice fed HFHS had significant reductions in the expression of upregulated genes seen in coronary arteries from WT mice fed HFHS diet (Sod1, Sod2, catalase, Cav-1, and iNos) and significant elevations in the expression of the downregulated gene (Pgc-1α). Interestingly, it has been reported that the loss of Pgc-1α is related to NO-to-H_2_O_2_ switch in flow mediated dilation in CAD patients [27, 28]. To determine if Pgc1α contributes to the NO-to-H_2_O_2_ switch in EDD in diabetes and if miR-21 regulates PGC1α, we isolated mCEC from WT and miR-21^-/-^ mice and treated the cells with HG. Figure 4D showed that compared to the WT, Pgc-1α expression was higher in miR-21^-/-^ mCEC subjected to both LG and HG treatment, suggesting that miR-21 deficiency upregulated Pgc-1α in mCEC. Each group had 3 animals and each animal were used for mCEC isolation separately. These data suggest that the deficiency of miR-21 reversed gene expression changes that contribute to vasodilation and ROS production in diet-induced diabetic mice.

### Diabetic mice have elevated levels of NO and H_2_O_2_ in the serum and elevated levels of superoxide in arteries

To understand the relationship between NO and H_2_O_2_ bioavailability and the relative contributions of NO and H_2_O_2_ during Ach-induced EDD in coronaries of diabetic mice, we assessed the production of NO and H_2_O_2_ in these mice. The level of NO was determined by measurement of NOx (Nitrite/Nitrate), which is highly correlated with NO level in serum [60]. Figures 6A and 6B showed that compared to WT mice, serum levels of NO and H_2_O_2_ were increased in db/db and diet-induced diabetic mice. Interestingly, the deficiency of miR-21 stopped this change in diet-induced diabetic mice. The gene expression in coronary arteries from diabetic animals and mCEC from WT and db/db mice (Figure 4.) showed the elevated expression of iNos and unchanged or decreased eNos, so the iNos might be the cause of elevated nitrate level. Figure 6C and 6D showed higher levels of ROS in the aortic arteries in db/db mice by staining with Dihydroethidium (DHE), a superoxide indicator, which is not an accurate index to show absolute amount of ROS but could show relative levels of superoxide in situ between the groups.

**Fig 6.**
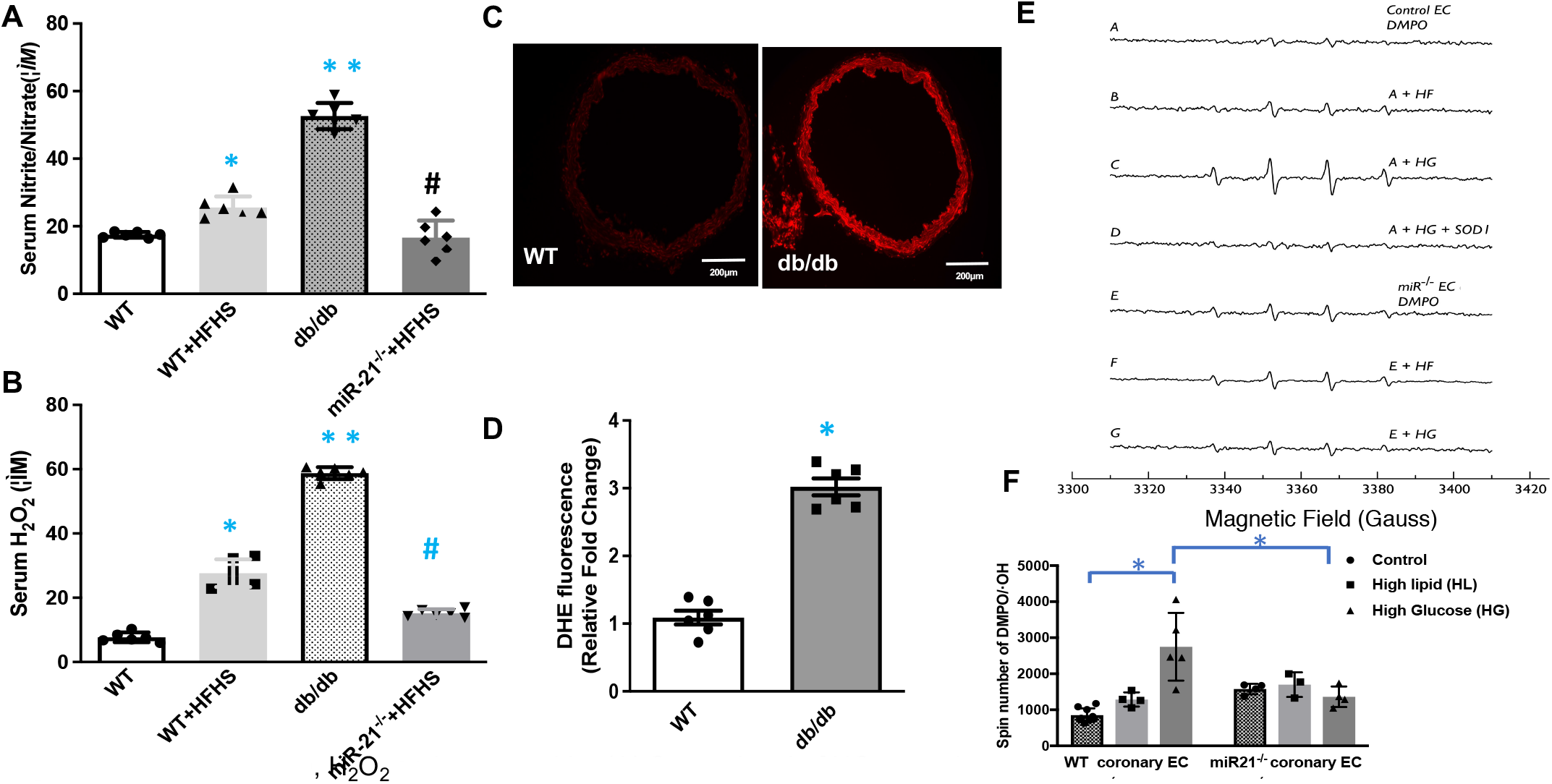
Assessment of NO, H_2_O_2_ and super oxide. (A) Serum level of NOx metabo +HFHS mice. (B) Serum level of H_2_O_2_ in WT, WT+HFHS, db/db and miR-21^-/-^ +HFHS mice. Compared to WT mice, level of NOx and H_2_O_2_ were increased and deficiency of miR-21^-/-^ reversed the changes in diet-induced diabetic mice. (A-B). (\**p*<0.05, \*\**p*<0.01 *vs.* WT, ^#^*p*<0.05 *vs.* WT+HFHS). (C) DHE staining in WT and DB aortic arteries (\**p*<0.05 *vs.* WT). n=6 mice/group. (D) The quantitation of C showing higher super oxide in arteries in db/db mice. (E-F) Superoxide detection in mouse coronary endothelial cells treated with high fat (HF, 150uM Linoleic Acid, Oleic Acid, and Cholesterol) and high glucose (HG, 25.5mM glucose) was performed using with DMPO (50 mM) spin-trapped and measured by EPR. Superoxide production under high sugar treatment increased significantly and knockout miR21 deficiency normalized the levels. (*p<0.05).

### miR-21 regulated ROS in mCEC

To study how miR-21 regulates ROS signaling in the endothelial cells of diabetes, mCEC were isolated and treated with HG and HL. Figure 6E and 6F showed EPR measurements of superoxide in mCEC under HG and HL treatment. WT mCEC subjected to HG showed significantly increased superoxide and the deficiency of miR-21 normalized the superoxide levels. Mechanistically, this may partially contribute to why miR-21 deletion prevented the switch from NO to H_2_O_2_ in EDD in diabetic coronary arteries.

### Implications of the NO-to-H_2_O_2_ switch in regulation of myocardial blood flow

As our previous results (Figures 1–4) have shown, EDD in coronary arteries under normal conditions is mediated by NO, but the mediator of EDD in coronary arteries is changed to H_2_O_2_ in diabetes. These results were observed *in vitro,* so whether there was a functional consequence of this switch with regards to myocardial blood flow (MBF) regulation *in vivo* is unknown. To determine if the NO-to-H_2_O_2_ switch has any effect on regulation of myocardial perfusion during metabolic hyperemia, MBF and hemodynamics were measured at baseline (during hexamethonium) and during norepinephrine (NE) administration in WT, db/db, and WT mice fed HFHS. Cardiac work (product of HR x MAP) during changes (i.v. norepinephrine) was calculated. HR and MAP are shown in Supplementary Figure 3. The relationship between DP and MBF is shown in Figure 7. The cardiac work is the double product of MAP and HR. After perfusion with NE to induce metabolic hyperemia, MAP and HR are increased. Blood flow in all groups increased when the double product increased; however, the hyperemic response (increase in blood flow) was less in the diabetics because the coupling of the metabolic demand and blood flow is impaired in diabetic mice. In db/db and diet-induced diabetic mice, MBF was significantly lower than MBF in WT mice during metabolic hyperemia produced by i.v. injection of NE (Figure 7A). L-NAME attenuated NE-induced hyperemia in WT mice (Figure 7B). However, Figure 7C and Figure 7D show that PEG-catalase, but not L-NAME, attenuated the NE-induced hyperemia in both db/db mice and diet-induced diabetic mice. Figure 7E–7G shows the MBF from Figure 7A-D. These results confirm that the mediator of coronary dilation in WT *in vivo* is NO-dependent, while the mediator of coronary dilation in diabetic mice *in vivo* is H_2_O_2_-dependent. These data confirm the occurrence of the NO-to-H_2_O_2_ switch *in vivo* in coronary dilation in diabetic animals. Moreover, compared to the WT mice, the myocardial blood flow in the diabetic mice is compromised; the switch of dilation mediator from NO to H_2_O_2_ may be a contributor.

**Fig 7.**
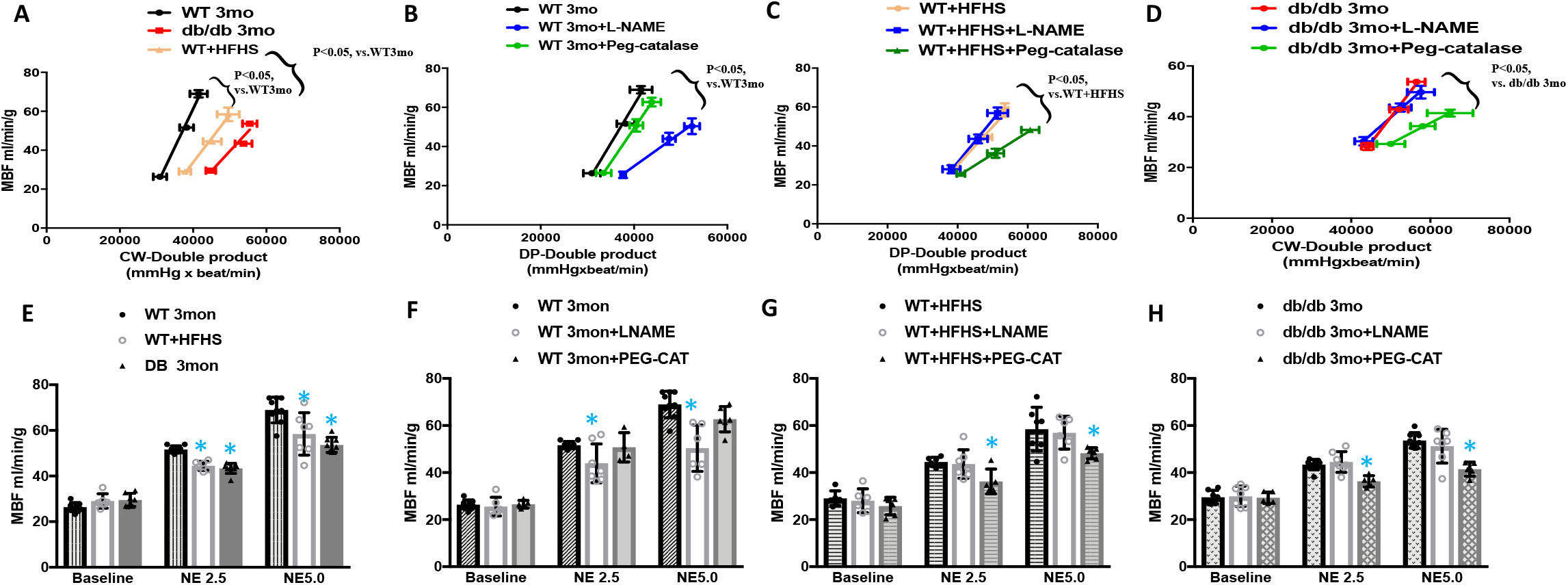
Relationship between Double Product (DP, DP=mean arterial pressure X heart rate) of cardiac work and myocardial blood flow (MBF). MBF and DP were measured (after hexamethonium) and during norepinephrine administration using contrast echocardiography and femoral artery catheterization. (A and E) in db/db or WT+HFHS mice, MBF was significantly lower than MBF in WT mice during metabolic hyperemia produced by i.v. injection of norepinephrine. (B and F) L-NAME attenuated the norepinephrine-induced hyperemia in WT mice, but Peg-catalase did not. (C-D,G and H) Peg-catalase, not L-NAME, attenuated the norepinephrine-induced hyperemia in db/db and WT+HFHS mice. This confirms that the mediator of coronary dilation in WT *in vivo* is NO dependent, while the mediator of coronary dilation in diabetic mice *in vivo* is H_2_O_2_ dependent. (\**P*<0.05-n=6 mice/group).

### miR-21 regulated PPARα and eNOS protein expression

Our gene expression data in coronary arteries and mCEC showed that miR-21 regulated PGC1α mRNA expression. PGC1α plays important roles in regulating microcirculation in CAD and obese patients [27, 28]. PPARα was predicted as an miR-21 target based on bioinformatics predictions (www.TargetScan.org) and it is reported that PPARα was downregulated in the retina of db/db mice and miR-21 inhibitor prevented the decreased PPARa [10]. miR-21 is reported to regulate eNOS in hypoxia and miR-21 mimic decreased eNOS expression [49]. To determine if miR-21 regulated the protein expression of the PPARα and eNOS, we performed western blot in miR-21^-/-^. Due to the size limitation of coronary arterioles and high expression of miR-21 in vessels, we performed the western blot in cardiac tissues. Figure 8A and B showed that deficiency of miR-21 increased PPARα and eNOS. Functional protein association networks by STRING (https://string-db.org, Figure 8C) showed the signaling pathways linking miR-21 targeting PGC1α and PPARα and how PGC1 α and PPARα regulates genes involved in ROS pathways that were shown via qPCR in Figure 4.

**Fig 8.**
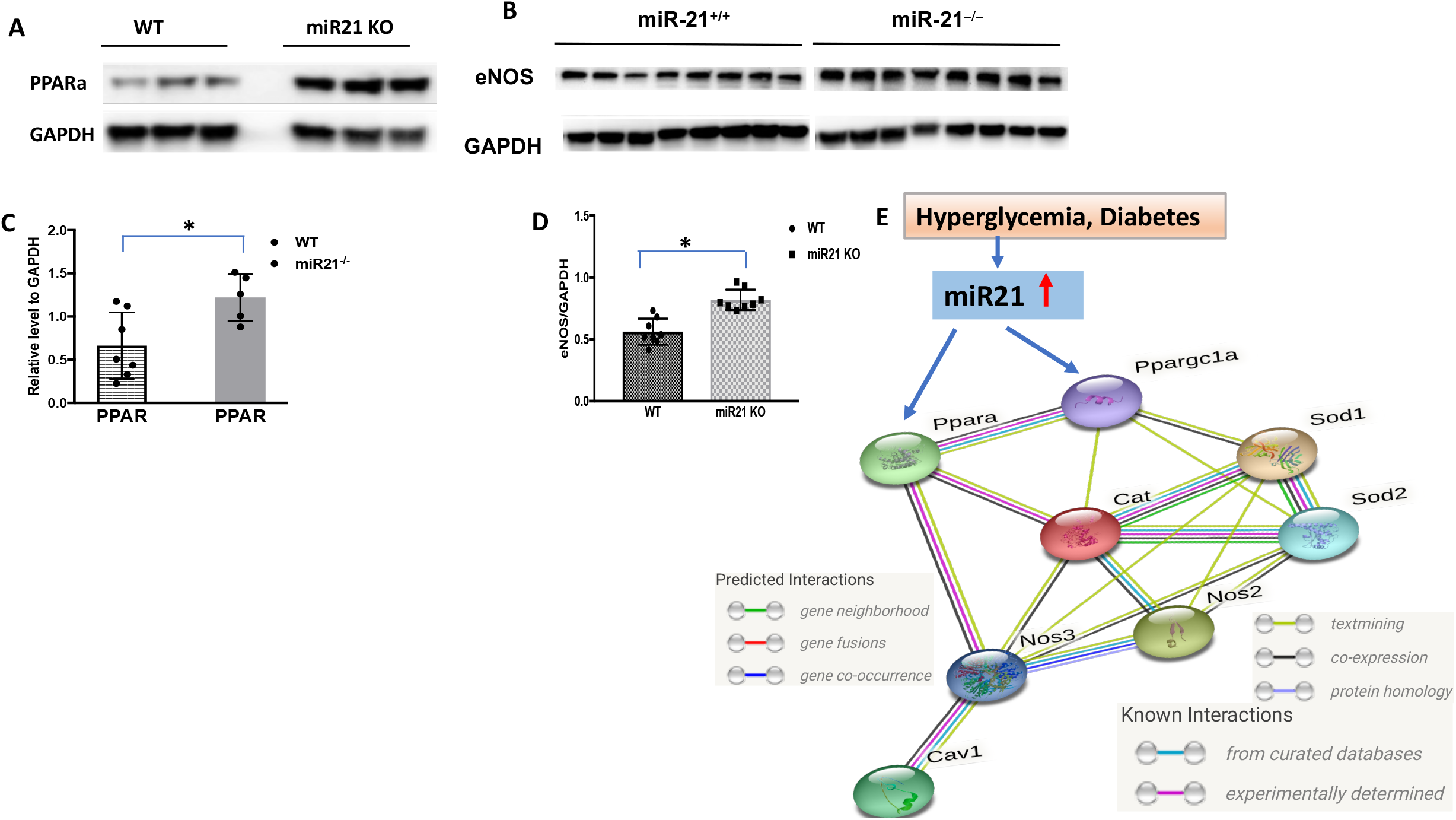
mi-R 21 regulates the PPARa and eNOS expression. (A-B). Western blot has shown that the deficiency of miR-21 increased PPARa and eNOS protein levels. (C-D) Quantitation of PPARa and eNOS by the relative level to GAPDH. (*p<0.05). (E) Functional protein association networks by STRING shows the signal pathway linking PGC1α and PPARa regulation of genes related to ROS homeostasis that were shown in qPCR in Figure 4.

## Discussion

Coronary microcirculatory function is important for the maintenance of proper myocardial perfusion, hence impairments in coronary dilation may lead to perfusion insufficiencies and myocardial ischemia. Coronary microvascular dysfunction in diabetes is well documented as an early event of diabetic cardiomyopathy [32], but the underlying mechanism is still an enigma. The current study provides some new insights into the coronary microcirculatory regulation in diabetes. Our results show for the first time that H_2_O_2_, not NO, as the mediator of acetylcholine – induced EDD in the coronary arteries from db/db as well as in diet-induced diabetic mice. In most cases, NO is the predominant mediator during Ach-induced EDD, so our data suggest that there is a diabetic switch from NO to H_2_O_2_ during acetylcholine-induced EDD in coronary arteries. This is the first report that recapitulates the clinical observation of this NO-to-H_2_O_2_ switch in flow-mediated EDD in CAD by Dr. Gutterman and colleagues. While senescence is usually linked to the endothelial dysfunction in diabetes [7], this study clearly showed that the NO-to-H_2_O_2_ switch during Ach-induced EDD in diabetic coronary arteries is not age-related. We have shown that NO is still the mediator of Ach-induced EDD in 32-month-old WT mice, while H_2_O_2_ is the mediator of Ach-induced EDD in 3-month-old db/db mice. These results are also consistent with reports from clinical studies in which an NO-to-H_2_O_2_ switch did not occur in elderly healthy patients [7]. Importantly, the deficiency of miR-21 was able to stop the NO-to-H_2_O_2_ switch during Ach-induced EDD in diet-induced diabetic mice. This suggests that miR-21 is an important regulator of the endothelial and coronary microvascular function in diabetes. Lastly, this is the first report that verified the NO-to-H_2_O_2_ meditator switch in coronary EDD *in vivo.* Most vasodilation studies rely on the *ex vivo* myograph or pressure gradient vasodilation. Our study assessed the reliance on specific vasodilators by i.v. injection of vasodilatory pathway inhibitors / disruptors (L-NAME and PEG-catalase) to investigate the effect of the NO-to-H_2_O_2_ switch on the regulation of MBF *in vivo.*

One aspect of our study that bears further consideration pertains to the models of type 2 diabetes to used. While various mouse models of diabetes exist, db/db mice are often used as a genetic means to recapitulate the diabetic phenotype, with these mice begin exhibiting a phenotype as early as 6 weeks of age. Although this appears to be completely unnatural and without parallel in humans, we note that type 2 diabetes is a rising threat in youth [2]. However, the db/db mouse has been criticized because of abnormal leptin signaling. Therefore, we also utilized a diet-induced diabetic mouse model that better reflects those patients who consume an unhealthy diet (high in fat and sugar) and progress to the development of type-2-diabetes [19, 46, 55]. Like these patients, our mouse model requires a longer period (5-6 months) to display a diabetic phenotype, especially with regards to metabolic derangements and cardiovascular dysfunction based on our preliminary data on mouse and rat.

We acknowledge the limitations associated with our current study and understand the importance of clinical / translational studies that ultimately promote improvements in patient treatment. However, mouse / cell models provide highly viable and accessible means to study mechanisms / pathways via gain and loss of function genetic manipulations that would be impossible for human studies. Novel disease targets discovered in basic cardiovascular research will eventually need to be tested in patients to promote the bench to bedside course of scientific discovery. One aspect of our study deserves emphasis; namely, our model of the switch from NO to H_2_O_2_ in mediating endothelial dependent vasodilation in diabetes mimics the finding in human patients with CAD [6, 21].

The balance of NO and H_2_O_2_ is important for maintaining endothelial homeostasis and function. Under normal conditions, NO is the major EDRF and contributes to the EDD of coronary arteries. Our results have shown that NO is the dominant mediator of acetylcholine -induced EDD in aortic arteries from WT and diabetic mice, which is consistent with the literature in that NO predominantly regulates the tone of large conduit vessels [39]. Our data also show that NO is the major and dominant mediator of acetylcholine -induced EDD in healthy coronary arteries, but not in diabetic coronary arteries. Previous reports have shown that L-NAME did not inhibit EDD in coronary arteries in db/db mice [4, 12, 48]. However, we are the first to report the major mediator of acetylcholine -induced EDD is H_2_O_2_ and identify the NO-to-H_2_O_2_ switch in diabetic coronary dilation. When NO bioavailability declines in diabetes, H_2_O_2_ compensates to mediate the EDD. Although, both NO and H_2_O_2_ are dilators, their inherent properties are very different [3, 62]. Endothelial and myocardial-derived H_2_O_2_ plays an important role in EDD in coronary circulation. H_2_O_2_ also plays important roles in pathological conditions such as atherosclerosis and hypertension with involvement during compensation of vasorelaxation in large vessels [33]. The adverse outcome of relying on H_2_O_2_ as a dilator in coronary circulation might be coronary microvascular dysfunction, which is the early event for both diabetic cardiomyopathy and coronary artery diseases. In that way, the NO-to-H_2_O_2_ switch in EDD could be a pathological indicator of coronary microvascular dysfunction in diabetes.

As to the underlying mechanism of the NO-to-H_2_O_2_ switch in Ach-induced EDD in diabetic coronaries, our study suggested that an imbalance of NO and H_2_O_2_ might be causal. Compared to WT mice, the serum levels of NO and H_2_O_2_ is increased in diabetic mice, which suggested a pathological increase in the production of NO and H_2_O_2_. Levels of NO and H_2_O_2_ in the coronary arteries are important for vasodilation, and especially for the NO-to-H_2_O_2_ switch in EDD in diabetes. Importantly, measuring NO in real-time is very challenging due to the short half-life (milliseconds) in biologic systems. Because NO is rapidly oxidized to nitrite and nitrate inorganic anions, they that have been considered as markers of NOS activity. The bioactivity of NO is partly regulated by its rapid oxidation to nitrite or nitrate, where nitrate is the predominant NO oxidation product in the circulation, but diet is the main contributor of exogenous nitrate. In blood and tissues, nitrite can be further reduced to nitric oxide and other bioactive nitrogen oxides [63]. Taken together, the measurement of NOx is an indirect but useful measurement NO. Our gene expression profiling also shows a change in the expression of genes that regulate ROS mitigation and NO bioavailability (eNos, iNos, Cav-1, catalase, Sod1 and Sod2) in diabetic mice [40, 53, 56]. Interestingly, Pgc-1α expression was significantly reduced in WT+HFHS and db/db mice. Our study using mCEC from WT and miR-21^-/-^ mice showed that Pgc1α expression was upregulated in miR-21^-/-^ cells under LG and HG compared to the WT. These data suggest PGC1α signaling to be involved in the NO-to-H_2_O_2_ switch in EDD in diabetic coronary arteries and miR-21 regulates PGC1 α signaling. PGC1 α is reported to have a host of functions including; antioxidant effects [23], prevention of oxidative stress [54], reduction of the expression of antioxidant-associated genes [11], reversed the overexpression of Sod1, Sod2 and catalase induced by ROS[16], diminished iNos as well as its NO by-product [18], promote eNOS expression [14] and is involved in regulating Cav-1 promoters [34]. Importantly, the loss of PGC-1α in non-CAD coronary arterioles produced a diseased (CAD) phenotype characterized by a shift from NO-to H_2_O_2_-mediated dilation [28]. This suggests that PGC-1α plays an important role in the NO-to-H_2_O_2_ switch by regulating oxidant / antioxidant gene expression in diabetic coronary arteries.

Recently, miR-21 has emerged as a big player in various cardiovascular diseases, where it can assume pathogenic and protective roles. Clinicals studies have used circulating miR-21 as a biomarker for disease [52, 58], but the lack of validation with regards to tissue specific measurements in patient samples have made the role of miR-21 in diabetes elusive. miR-21 plays important roles in metabolism [66],[9, 51], with implications in diabetic pathology. miR-21 is enriched in EC, where it regulates NO production, and ROS-homeostasis by targeting SOD2. miR-21 has also shown to influence cell proliferation and apoptosis [31, 61, 67], and modulated vascular disease and remodeling [57, 59]. Additionally, miR-21 was also found to be up-regulated by replicative and stress-induced senescence [17]. Although our data suggest the NO-to-H_2_O_2_ switch was not age related, this alludes to the complexities of diabetes and the need for contextdependent interrogation of miR-21 and its functions.

Interestingly, miR-21 has shown to be protective during diabetic cardiomyopathy-induced diastolic dysfunction [15], and confers vascular protection during ischemic preconditioning [64]. However, miR-21 antagomir also enhances the impaired process coronary collateral growth in rat model of metabolic syndrome [25],[24], and deficiency of miR-21 improves the outcome of diabetic retinopathy [10]. Because systemic delivery of miR-21 antagomirs lacks tissue specific targeting, we could not exclude the effect of miR-21 antagomirs on the other organs. To determine miR-21 and other gene expression during the process of coronary collateral growth in diabetic animals, we performed RNA-sequencing analysis. Our unpublished data showed that miR-21 was one of the top genes that was upregulated in the collateral dependent zone of Zucker Obese rat hearts during the collateral growth time window. This suggested that miR-21 is an important regulator vascular endothelial function. In this study, the deficiency of miR-21 stopped the NO-to-H_2_O_2_ switch during Ach-induced EDD in diet-induced diabetic mouse coronaries. Ultimately, our data suggested that miR-21 holds an important role for the regulation of the endothelial and coronary microvascular function in diabetes.

In this study, we explored how miR-21 regulates the NO-to-H_2_O_2_-switch in coronary arteries of diabetic mice. Our data suggest that miR-21 directly targets PGC1α and PPARα which are important regulators of ROS signaling, metabolism and diabetes. PGC1 α has shown to play important roles in microcirculation and is involved in the switch of NO-to-H_2_O_2_ during EDD in CAD patients. Functional protein association networks by STRING revealed PPARα interaction with PGC1α, and the regulation of eNOS and Catalase. PGC1α may also impart regulation on SOD1 and SOD2, while upstream signaling of eNOS and Catalase regulates CAV-1, iNOS and eNOS, with interaction between iNOS, SOD1, and SOD2. Our gene expression data in coronaries confirm the signaling network generated in STRING to further drive our hypothesis that miR-21 contributes towards diabetic microvascular dysfunction.

Therefore, miR-21 may represent a novel therapeutic target for the treatment of coronary microvascular diseases in diabetes. Importantly, use of miR-21 antagomirs will be feasible for clinical studies to influence the pathological switch of NO to H_2_O_2_ and thereby improve the coronary microvascular function and possibly prevent the progression to diabetic cardiomyopathy. miR-21 has been proposed as a therapeutic target in hypertension and heart failure [5, 35], and anti-miR-21 oligonucleotides have been used as inhibitors for studies of heart failure and hypertension. RG-012, is a relevant example of a miR-21 inhibitor that is used in patients with kidney fibrosis[22], but the confounding nature of miR-21 in the context of various disease requires careful scrutiny so that off-targeted effects are limited. Systemic delivery lacks tissue specific targets and attempts to localize miR-21 inhibition may be of interest, particularly in cases of overlapping disease where the roles may be opposing.

We acknowledge the limitations associated with our current study and understand the importance of clinical / translational studies that ultimately promote improvements in patient treatment. However, mouse / cell models provide highly viable and accessible means to study mechanisms / pathways via gain and loss of function genetic manipulations that would be impossible for human studies. Novel disease targets discovered in basic cardiovascular research will eventually need to be tested in patients to promote the bench to bedside course of scientific discovery. While the use of human coronary EC purchased from Lonza and more patient samples surely would have improved the study, acquiring clinical specimens is challenging, especially obtaining live cardiac tissue. Several different isolated vessel techniques have been used to further our understanding of vasomotor regulation. In the current study, we utilize the agonist acetylcholine to induce vasodilation because this method puts emphasis on the NO, which is central to endothelial dysfunction. Another method known as flow-mediated-dilation (FMD), interrogates vasomotor reactivity during changes in pressure gradients that produce flow to elicit vasodilation via shear responsive elements on the endothelium through PG and NO [42]. FMD may be more physiological, but difficulties arise when dealing with mouse coronaries, which are highly branched (unlike aortic or mesenteric arteries) and prove challenging to study via FMD.

In the current study, we investigate the coronary vasodilation by acetylcholine-induced EDD via myography, which is widely used in literature. In addition, we do realize the dosage of acetylcholine might not be physiological, but clinical studies utilized acetylcholine as an endothelium-dependent probe to measure the coronary blood flow reserve in patients with the dosage range from 10^-6^ to 10^-4^ uM at infusion rate 48-120 mL/h for 3 minutes infusion duration and infused dosage from 0.364 μg to 108 μg [12]. The concentration of acetylcholine in our studies was 10^-9^ to 10^-5^ μM and is like the dosage in clinical studies. Furthermore, to study the NO-to-H_2_O_2_ switch under physiological conditions, we performed an *in vivo* study to test the effect of L-NAME and PEG-catalase on the myocardial blood flow in diabetic mice and WT mice. Our *in vivo* data confirm that the endothelial dependent dilator switches from NO to H_2_O_2_ in the diabetic coronary circulation observed *ex vivo* occurs *in vivo* in diabetic mice.

In summary, the present study reports that the mediator in Ach-induced EDD switches from NO to H_2_O_2_ in coronary arteries of db/db mice and diet-induced diabetic mice. This is the first mouse model that recapitulates the clinical observation of the NO-to-H_2_O_2_ switch in EDD in CAD patients. More importantly, our study shows this switch also occurs *in vivo* in diabetic mice, thereby compromising their myocardial blood flow. Deficiency of miR-21 could prevent the NO-to-H_2_O_2_ switch in diet-induced diabetic mice, which suggests that miR-21 might be a new therapeutic target in coronary microvascular diseases and microvascular dysfunction in diabetes.

## Acknowledgements

This research is funded by National Institutes of Health grant 2R01HL103227-05 (YZ, LY), 1R01HL135110-01 (WMC, LY), 1 R01 HL137008-01A1 (LY)

